# Noninvasive Prenatal Screening at Low Fetal Fraction: Comparing Whole-Genome Sequencing and Single-Nucleotide Polymorphism Methods

**DOI:** 10.1101/096024

**Authors:** Carlo G. Artieri, Carrie Haverty, Eric A. Evans, James D. Goldberg, Imran S. Haque, Yuval Yaron, Dale Muzzey

## Abstract

**Objective:** Performance of noninvasive prenatal screening (NIPS) methodologies when applied to low fetal fraction samples is not well established. The single-nucleotide polymorphism (SNP) method fails samples below a predetermined fetal fraction threshold, whereas some laboratories employing the whole-genome sequencing (WGS) method report aneuploidy calls for all samples. Here, the performance of the two methods was compared to determine which approach actually detects more fetal aneuploidies.

**Methods:** Computational models were parameterized with up-to-date published data and used to compare the performance of the two methods at calling common fetal trisomies (T21, T18, T13) at low fetal fractions. Furthermore, clinical experience data were reviewed to determine aneuploidy detection rates based on compliance with recent invasive screening recommendations.

**Results:** The SNP method’s performance is dependent on the origin of the trisomy, and is lowest for the most common trisomies (maternal M1 nondisjunction). Consequently, the SNP method cannot maintain acceptable performance at fetal fractions below ~3%. In contrast, the WGS method maintains high specificity independent of fetal fraction and has >80% sensitivity for trisomies in low fetal fraction samples.

**Conclusion:** The WGS method will detect more aneuploidies below the fetal fraction threshold at which many labs issue a no-call result, avoiding unnecessary invasive procedures.

## BULLETED STATEMENTS: What’s already known about this topic?

- The two most popular noninvasive prenatal screening (NIPS) methodologies, the single nucleotide polymorphism (SNP) and the whole-genome sequencing (WGS) methods, report comparable performance.
- However, failure rates vary by an order of magnitude between methodologies.
- A large component of failure is insufficient fetal fraction, creating “no-call” test results.
- The American Congress of Obstetricians and Gynecologists (ACOG) indicates that reported sensitivity is often inflated due to exclusion of failed samples from calculations.

## What does this study add?

- Unlike the WGS method, the SNP method is sensitive to the origin of aneuploidy, and performs poorly at low fetal fraction on the most common fetal trisomy (maternal M1 nondisjunction).
- Offering invasive testing to all cases of “no-call” results will likely increase the rate of procedure-related loss.
- The WGS method maintains high specificity and can detect a higher proportion of aneuploidies in low fetal fraction samples without unnecessary invasive tests.

## Introduction

Noninvasive prenatal screening (NIPS) for fetal aneuploidies using cell-free DNA (cfDNA) has been widely adopted in clinical practice due to its improved accuracy as compared to traditional screening approaches (1). Consequently, both the American Congress of Obstetricians and Gynecologists (ACOG) and the American College of Medical Genetics and Genomics (ACMG) recommend NIPS as a routine screening option (2,3).

Such performance improvements have been enabled by developments in next-generation sequencing (NGS) technologies, which are employed by most clinical NIPS laboratories. NGS involves the generation of millions of short sequences (“reads”), each originating from a specific chromosomal segment(4), that provide information about both the genotype and relative abundance of the site in the genome (e.g., the X chromosome receives more reads in XX females than in XY males). The two most widely offered NIPS methodologies differ based on which data they use to detect fetal aneuploidies.

The single-nucleotide polymorphism approach (“SNP method”), measures the relative proportion of maternal and fetal genotypes among cfDNA fragments, and tests whether the observed patterns on specific chromosomes are more consistent with disomic or aneuploid fetal expectations (5). Alternatively, the whole-genome sequencing approach (“WGS method”), measures the relative abundance of cfDNA from whole chromosomes in the maternal blood, testing whether certain chromosomes show elevated or reduced numbers of reads, consistent with fetal aneuploidies (6). Despite differences in the underlying signals, meta-analyses have found that both approaches share comparable clinical sensitivities for detecting common aneuploidies: trisomy 21 (Down Syndrome, T21), trisomy 18 (Edwards Syndrome, T18), trisomy 13 (Patau Syndrome, T13), and monosomy X (7,8).

A key concern has been how these methods perform on the minority of patients with low abundance of circulating fetal DNA (i.e., low fetal fraction samples), where the threshold between signal and noise blurs (9). Low fetal fraction is associated with high maternal body-mass index and certain fetal aneuploidies (1,10). In order to maintain high per-patient sensitivity, some tests avoid reporting results to patients with fetal fractions below a preset threshold - referred to as a “no-call” result. Patients who receive a “no-call” may submit a second blood draw or be offered invasive testing as higher rates of aneuploidy have been reported in such samples (11).

However, it is not clear that the strategy of combining “no-calling” with invasive diagnostic follow-up improves the detection rate of fetal aneuploidies as compared to simply calling low fetal fraction samples with reduced sensitivity: clinical experience shows that only 56.5% of patients submit a second sample and at least 25% of these fail again due to low fetal fraction (10,11). Consequently, both ACOG and ACMG recommend that patients receiving a “no-call” be offered invasive diagnostic testing and that a second blood draw is not appropriate (2,3). Yet compliance with this recommendation is far from perfect: even among women who screen positive for aneuploidies using either NIPS or conventional screening methods, only ˜55% seek confirmatory invasive testing (10,12). Finally, an additional consideration is that invasive tests cause procedure-related pregnancy loss in approximately one in 500 cases, which could affect a substantial number of patients in the context of routine NIPS (13).

In this study we use up-to-date published validation reports to model the performance of the WGS and SNP methods at low fetal fractions in order to determine whether returning a “no-call” result leads to a higher rate of detection of common aneuploidies as opposed to providing results for all cases regardless of fetal fraction.

## Methods

### Simulating the WGS method

A detailed description of the equations and procedures underlying both methods is available in the Supplementary Material accompanying this manuscript and all code used to generate and analyze the data is available at https://github.com/counsylresearch/artieri_et_al_nips_at_low_ff. Here we briefly outline the key references and assumptions used to assess their performance. The WGS method was simulated according to the parameters in Jensen et al. (14): 16 million reads per sample counted in 50 kb bins along chromosomes 21, 18, and 13. Chromosomal sizes, (GRCh37/hg19, Feb. 2009) (15) were reduced by 10% to account for exclusion of poorly-performing bins (resulting from high-GC content or the presence of repetitive elements) (14). As proper normalization leads to bin counts being distributed according to Poisson expectations (Supplemental Figure S1) (16), bin depths were sampled from a Poisson distribution with mean based on chromosomal ploidy and fetal fraction (see Supplemental Methods).

Sample-specific z-scores for each chromosome were generated by simulating batches of 100 samples (as could be run on a single Illumina v4 High Output sequencing flowcell on a HiSeq 2500 instrument [Illumina, San Diego, CA, USA]), randomly drawn from a population with trisomy prevalence of 3.3%, T21; 1.5%, T18, 0.5%, T13 (8). A single z-score from each of a disomic and trisomic sample (if present) were chosen at random from each batch until 10,000 of each were sampled at a given fetal fraction. Samples were called trisomic if their z-score was greater or equal to three. Sensitivity was calculated as the fraction of trisomic samples correctly called, while specificity was the fraction of disomic samples not called trisomic.

### Simulating the SNP method

For the SNP method, parameters were obtained from Ryan et al. (11). 13,392 SNP sites were equally divided among chromosomes 21, 18, 13, and X. The mean number of NGS reads-per-SNP, 859, was obtained by dividing 11.5 million, the average sequencing depth of samples with <7% fetal fraction, by the total number of SNPs. As the variance in counts per SNP and allelic balance are not published, we modeled these parameters to generate data consistent with published figures (20) and show that all conclusions drawn are robust to the specific parameter values (Supplemental Figures S2–5). Samples were simulated in accordance with allele distributions as expected based on the parent- and meiotic-stage-of-origin of the trisomy and classified according to the approach outlined in Rabinowitz et al. (17) (see Supplemental Material).

Sensitivity of the SNP method at each fetal fraction was calculated by simulating 10,000 samples for each of the four types of meiotic nondisjunction and determining the proportion of trisomic samples for which the trisomy hypothesis had log-odds ratio (LOR) below the threshold at which 99.87% of disomic samples would be called disomic (corresponding to the same specificity as the WGS test). The aggregate sensitivity of the SNP method was determined by calculating the weighted expectation of the sensitivity of detection of each fetal ploidy hypothesis multiplied by its prevalence (18).

### Determining clinical outcomes

The proportion of patients who submit redraws, 56.5%, was obtained from Dar et al. (10). We used the highest reported probability of obtaining a successful result upon redraw for samples below 3% fetal fraction: 74% (11). As it is unlikely that “no-calls” lead to termination of pregnancy, we used the proportion of remaining patients receiving a positive NIPS call in Dar et al. (10) who elected invasive testing, 55%, to estimate the maximal rate at which patients receiving a “no-call” would seek invasive testing. A similar value, 57%, was reported for conventional first trimester screening (12). The rate of procedure-related loss, 0.002, was obtained from Yaron (19).

To calculate the detection sensitivity for the WGS method for samples ≤ 2.8% fetal fraction (11), we determined the frequency of observing samples in bins of 0.1% fetal fraction from 0 to 2.7% by fitting a beta distribution to the parameters reported in Nicolaides et al. (19) (median: 0.1; 25/75 percentiles: 0.078 and 0.13) using the ‘optimize.brute’ function in SciPy beginning with priors of zero and minimizing the sum of squares deviation from the 25th and 75th percentiles (20). We then obtained the dot product of the sensitivity calculated at each fetal fraction bin multiplied by its relative prevalence among samples in the 0 to 2.7% range.

### Statistics and Data

All statistics were calculated using SciPy in Python (version 0.17) (20). The output of all analyses as well as the Python code required to reproduce all results is available under the Creative Commons Attribution-NonCommercial 4.0 International Public License at https://github.com/counsylresearch/artieri_et_al_nips_at_low_ff.

## Results

### Comparing NIPS methods

To compare the WGS and SNP methods, we developed a computational framework with two steps: (1) simulation models that mimic the raw data generated by each method, and (2) aneuploidy-calling algorithms that process the simulated data and yield ploidy calls for trisomies in chromosomes 21, 18, and 13 (see Methods). The simulations allowed us to model an arbitrary number of pregnancies over a precise range of fetal fractions and calculate analytical performance in terms of both sensitivity and specificity. To ensure a fair comparison between methods, simulation parameters and calling algorithms were drawn directly from the most up-to-date published reports, such as peer-reviewed manuscripts and patents (see Methods).

The WGS method partitions each chromosome into equal sized bins and tallies the number of reads per bin, normalized for GC content and repetitive regions (14). Trisomies manifest as a higher number of reads per bin relative to the disomic background, with read excess proportional to the fetal fraction (e.g., 55 reads in trisomic chr21 bin vs 50 reads in disomic chr3, Figure 1, top). In contrast, the SNP method measures relative counts among alleles at pre-selected, polymorphic sites on the chromosomes of interest. Trisomies manifest as a global shift in allelic counts relative to disomic chromosomes (Figure 1, bottom).

**Figure 1.**
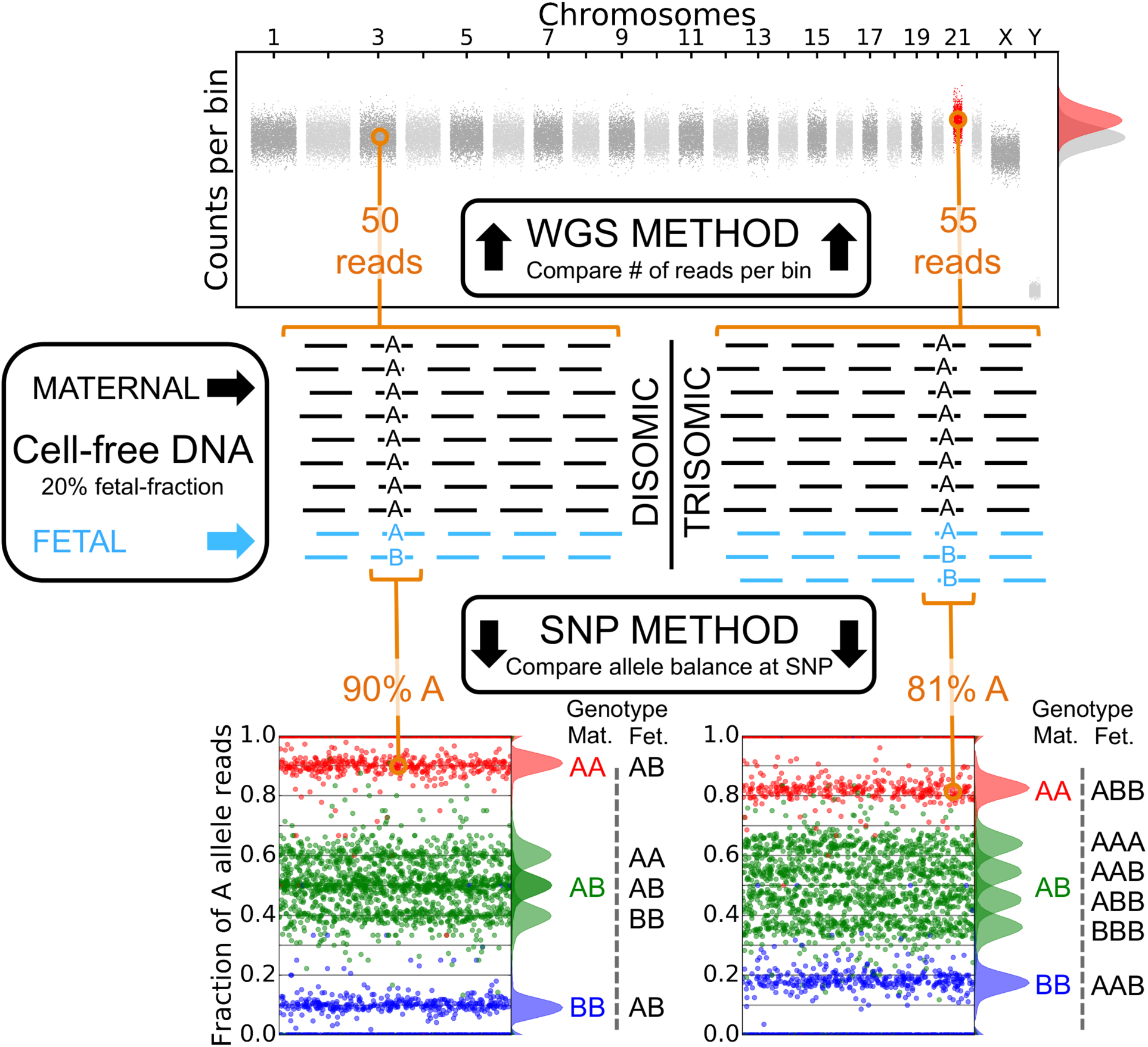
Overview of the WGS and SNP methods. Both methods take advantage of NGS reads originating from the mixture of maternal and fetal cfDNA (in this example, fetal cfDNA constitutes 20% of the pool). The WGS method (top) divides the genome into equally-sized bins and counts the number of reads mapped to each bin. As illustrated, the presence of a fetus affected with trisomy 21 leads to an increase in the distribution of counts-per-bin originating from chromosome 21 (55 reads, red) relative to the euploid background (50 reads, grey). Detection of this increase forms the basis of the test. Alternatively, the SNP method (bottom) measures the relative abundance of alleles at polymorphic sites in the cfDNA. Fetal aneuploidies lead to predictable shifts in the frequency of allelic counts based on the possible combinations of maternal and fetal genotypes (see below). By aggregating the signal across many SNPs on a given chromosome, the algorithm calculates a likelihood that the overall pattern of allele frequencies is more consistent with a normal versus an aneuploid fetus. The strength of the aneuploid signal in either method is proportional to the fetal fraction, and so too is the sensitivity of detection. Note that the number of reads illustrated in the figure are substantially lower than those published in validation reports (see Methods).

### Influence of parent-of-origin on SNP and WGS methods

The expected deviation in allele frequencies caused by aneuploidies in the SNP method differs based on the parental- and meiotic-stage of origin of the aneuploidy (Figure 2; Supplemental Figure S6), the rates of which vary substantially: 70% of nondisjunction events occur during maternal meiosis phase I (M1), leading to fetal inheritance of both maternal chromosomes, while 20% occur during meiosis phase II (M2), causing fetal inheritance of two copies of a single maternal chromosome (21). The remaining nondisjunctions are paternal in origin, with 3% originating from M1, and 7% from M2 (18) (Figure 2A).

**Figure 2.**
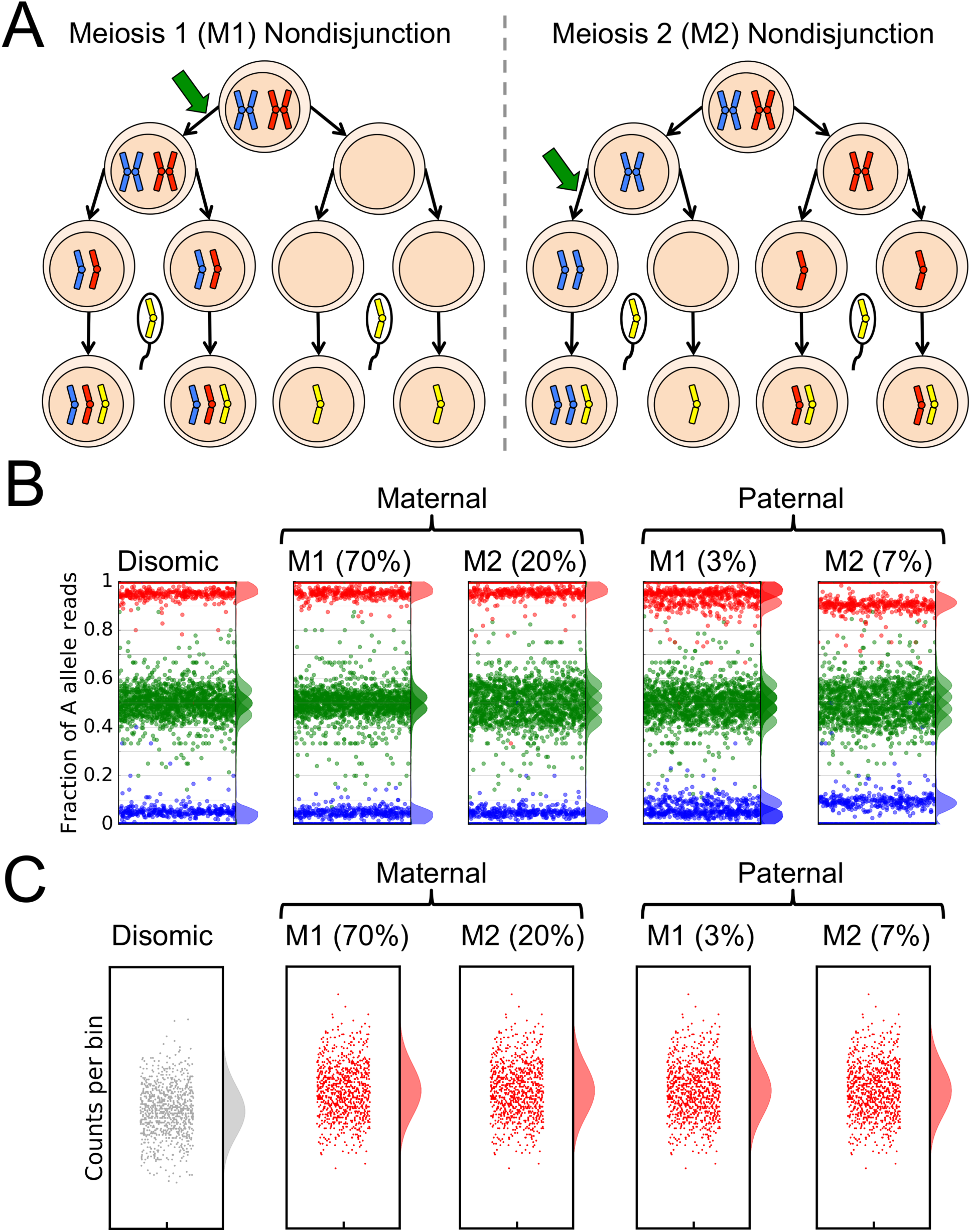
The parent-of-origin and meiotic-stage-of-origin of the aneuploidy produce different expected distributions of allele-frequencies with the SNP method. A) Illustration of the two origins of maternal trisomies. Trisomies can originate either from meiosis stage 1 (M1; left) or stage 2 (M2; right) nondisjunction. In M1 nondisjunctions, the oocyte inherits one copy of each of the different maternal chromosomes (marked in blue and red). Upon fertilization, the paternal chromosome (yellow) is added. In such cases, SNP-based analysis for each locus will yield one of six possibilities, resulting from the admixture of maternal and fetal cfDNA (AA|A, AA|B, AB|A, AB|B, BB|A, and BB|B, where the maternally inherited genotype is to the left and the paternal is to the right). In M2 nondisjunctions, the oocyte inherits two copies of the same maternal chromosome, leading to only four possibilities (AA|A, AA|B, BB|A, and BB|B). Cases of paternal nondisjunctions follow the same lines of reasoning with the parental origins reversed. Adapted from Karp (34). B) The four possible parental origins of trisomies produce distinct signals with the SNP method. Simulated allele-frequency distributions for samples with 10% fetal fraction for each of the different origins of trisomies are shown along with their relative frequencies. The density plots to the right of each panel indicate the shape of the distributions (each corresponding to the admixture of maternal and fetal cfDNA). Note that the shifts in heterozygous fetal SNPs on a homozygous maternal background (blue and red dots above) are significantly more pronounced in the rare paternally-derived aneuploidies in comparison to the much more common maternally-derived aneuploidies when compared to a euploid sample. The allelic basis of all possible maternal-fetal cfDNA genotypic combinations is shown in Supplemental Fig S6. C) With the WGS method, all four trisomic origins lead to the same increase in the number of reads per bin for the trisomic chromosome.

In the SNP method, paternally-derived trisomies produce a much stronger signal than do the more common maternal trisomies, illustrated by the shift in the red and blue distributions in Figure 2B. Because the majority of allelic counts in cfDNA are maternal in origin, paternally inherited trisomies nearly double the presence of paternal-specific alleles, whereas maternally-derived trisomies only slightly increase the signal of one of the two maternal alleles (˜4%) (Table 1).

**Table 1.**
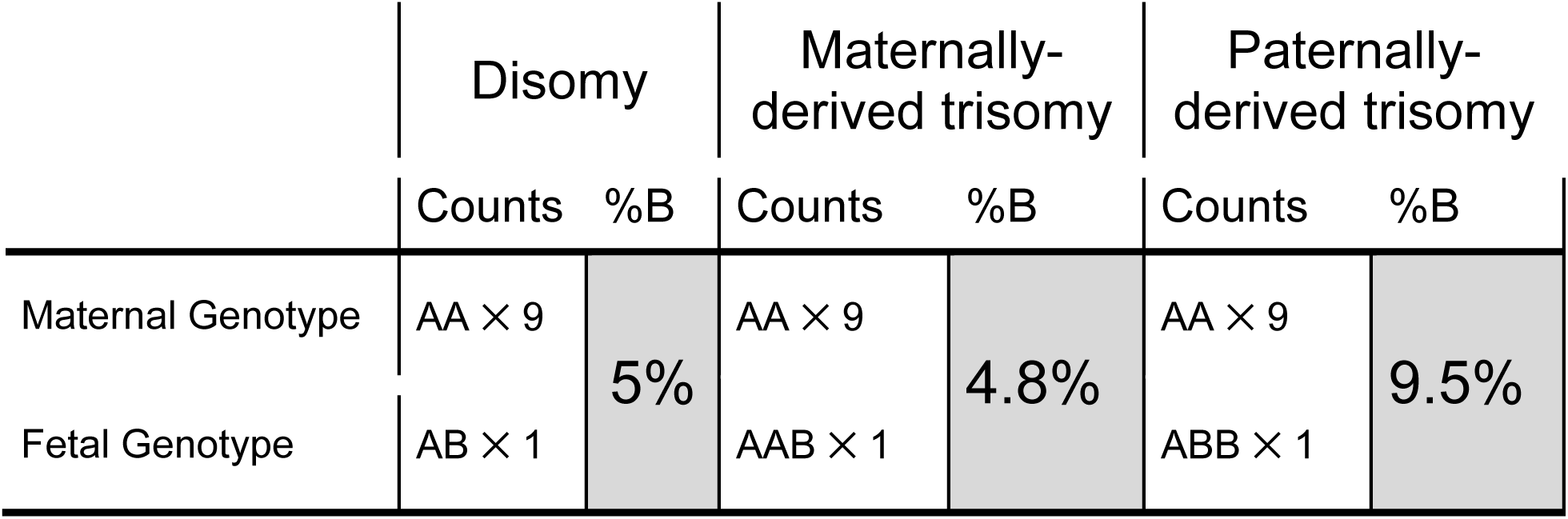
Rare paternally inherited trisomies produce a stronger SNP count signal than do more common maternally inherited trisomies. Values are shown for a 10% fetal fraction, a maternal genotype, AA, and a fetal genotype, AB. Paternal trisomies will increase the frequency of the paternally inherited B allele almost two-fold over euploid expectations, while the increase of the frequency of the A allele only shifts the expected abundance of the B allele by 0.2%.

Conversely, in the WGS method, all four origins of trisomy produce the same signal: an elevated number of NGS reads mapping to the trisomic chromosome (Figure 2C). Consequently, the sensitivity of the method is independent of the origin.

### Specificity and sensitivity of both methods as a function of fetal fraction

We compared the performance of the two methods at low fetal fraction by simulating 10,000 samples of each possible origin of trisomy for chromosomes 21, 18, and 13 at fetal fractions ranging from 0.1% to 4% (see Methods). Per-sample sequencing depths were obtained from published validation reports (11,14).

We determined calling performance by generating receiver-operating characteristic (ROC) curves and calculating the area under the curve (AUC) for each method as a function of fetal fraction (Figure 3A,B) (22). A test with high sensitivity and specificity will have AUC near 1. The WGS method processes ˜100 samples per batch and, for each sample and chromosome, assigns a z-score indicating the extent of which the counts in this sample deviate from the distribution of all other samples in the batch (23). Because the expected distribution of disomic samples is independent of fetal fraction, its specificity is a function of the z-score threshold (z ≥ 3) (9). This is confirmed by the AUCs, which are greater than 0.99 at fetal fractions above 1.5% for all three common trisomies (Figure 3A).

**Figure 3.**
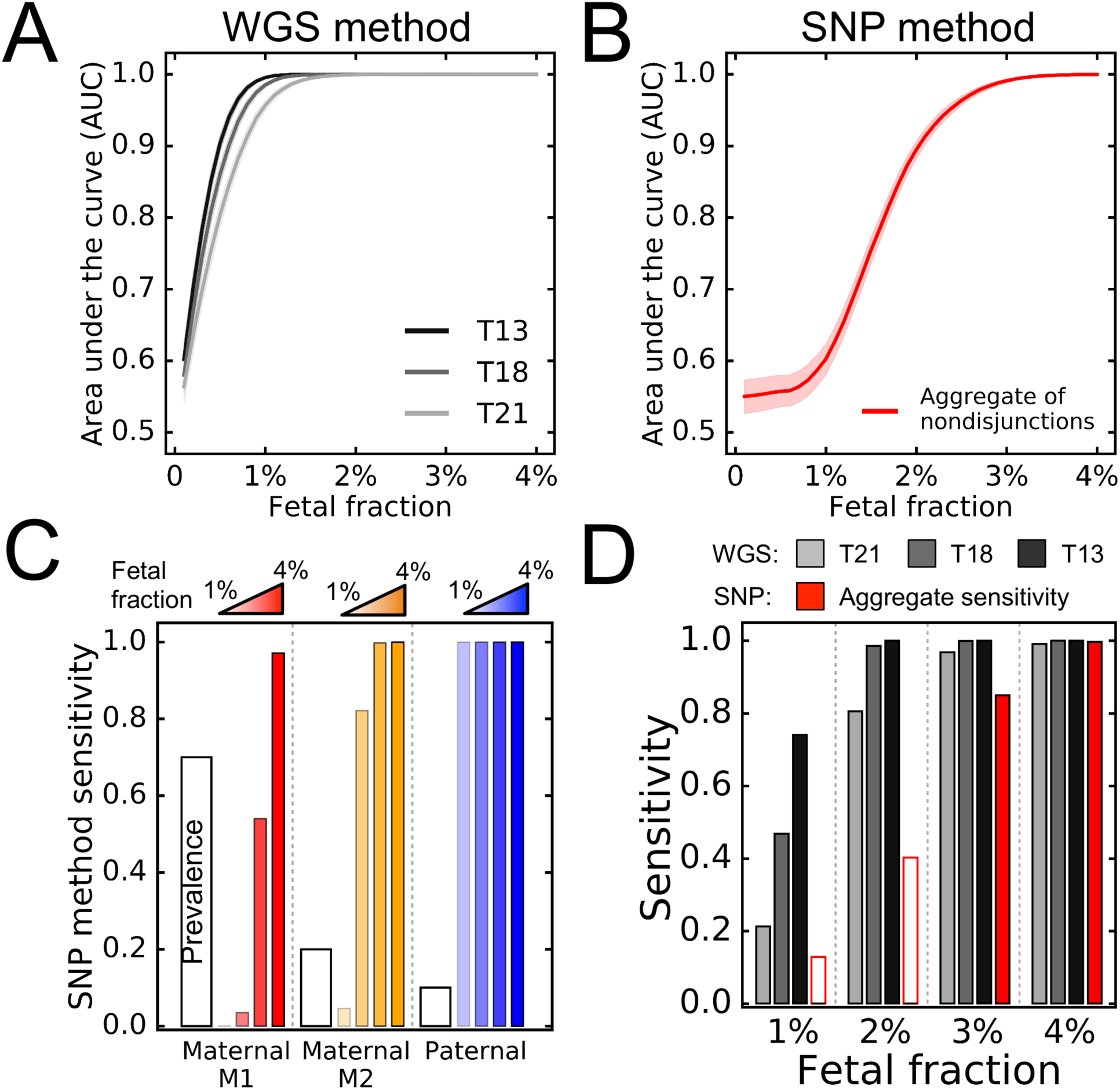
Comparison of performance characteristics of the two methods at low fetal fractions. Area-under the curve (AUC) values as a function of fetal fraction for the WGS (A) and SNP method (B) calling algorithms. The AUC captures the overall performance of disease classification and represents the probability that a random disomic fetal sample would have a higher z-score than a random aneuploid sample in the WGS method (or a higher likelihood of aneuploidy in the SNP method) (22,35). The WGS method achieves AUC > 95% for all three common trisomies at > 1 % fetal fraction. In contrast, the SNP method only achieves AUC ≥ 95% at fetal fractions ≥ 2.4%. The shaded areas above and below the curves represent the 95% confidence intervals on the estimates of AUC. C) Sensitivity of the SNP method at 99% specificity with respect to each of the trisomic origins (as both paternal non-disjunctions have identical sensitivity in this range, they are combined into a single category). The prevalence of each of the origins is shown as the white bar, while the colored bars indicate increasing fetal fraction in 1% intervals. Note that the sensitivity of the maternal M1 trisomy at 1% fetal fraction is zero, thus the leftmost bar is missing. Importantly, sensitivity is lowest for the most prevalent trisomies (maternal M1, 70%). D) Comparison of the sensitivities of the WGS method for each of the three common autosomal trisomies (grey bars) to the aggregate sensitivity of the SNP method (solid red bars). The aggregate sensitivity of the SNP method was obtained by summing the sensitivities scaled by prevalence across the three categories of trisomic origin. As illustrated by the simulations, the sensitivity of the WGS method improves with increasing chromosomal size. The aggregate sensitivity of the SNP method is shown as hollow bars at 1% and 2% fetal fraction as these are below the threshold at which all samples are “no-called”. In panels C and D, groups of bars are separated by dotted lines as a visual aid.

In contrast, the SNP method calculates a likelihood that each chromosome is disomic or trisomic on a per-sample basis, returning a log-odds ratio (LOR) where disomic samples have positive LOR and trisomic samples have negative LOR. The SNP algorithm shows poor differentiation between disomic and trisomic LORs at low fetal fractions as an AUC of 0.99 is not achieved below fetal fractions of 3% (Figure 3B). Setting a fetal fraction threshold below which all samples are “no-called” is thus appropriate to maintain SNP-method performance (e.g., ≤ 2.8% 11).

To translate AUCs into sensitivities, we calculated the maximum attainable sensitivity of the SNP method at each fetal fraction while maintaining specificity ≥ 99.87% (z ≥ 3 in the WGS method, see Methods). As expected, rare paternal trisomies could, in principle, be confidently detected at extremely low fetal fractions(Figure 3C). However, the most common fetal trisomies, maternal M1 nondisjunctions(70%), show the lowest sensitivity, with maternal M2 nondisjunctions(20%) falling in between the two extremes. We obtained the aggregate sensitivity of the SNP method at each fetal fraction by taking a prevalence-weighted sum of the sensitivities for each trisomie origin.

The sensitivity of the WGS method is dependent on the number of bins used to count reads and increases with chromosome size (Figure 1A) (16). Therefore, we calculated the sensitivity of the WGS method for each of the three common trisomies separately. In all cases, the SNP method shows lower aggregate sensitivity than the WGS method at low fetal fractions (Figure 3D).

### Clinical outcomes of low fetal fraction samples

Patients receiving a “no-call” have the option of sample redraw or invasive testing, however noncompliance may lead to undiagnosed aneuploidies. Therefore we assessed the T21 detection rate under two scenarios: (1) “no-calling” all samples below a fetal fraction of 2.8% (11), and (2) calling all samples using the sensitivity parameters established from simulations of the WGS method. We first calculated the sensitivity of the WGS method for all samples below 2.8% fetal fraction by summing the prevalence-weighted sensitivity at each fetal fraction (Supplemental Figure S7) (24). Applied to the simulated data, the WGS method shows a sensitivity of 86% for samples with fetal fraction < 2.8%.

Among patients initially receiving a “no-call” by the SNP method, approximately 42% will submit a redraw and receive a result (Figure 4A) (10,11). Subsequently, the rate of invasive procedures among remaining patients will determine the proportion of aneuploidies that are detected (we assume that redraws and invasive testing are 100% sensitive to establish an upper limit on the detection rate). The invasive test rate among patients receiving a “no-call” would have to exceed 76% to equal the sensitivity of the WGS method, which is unlikely given the upper-limit estimate of 55% (Figure 4B; see methods).

**Figure 4.**
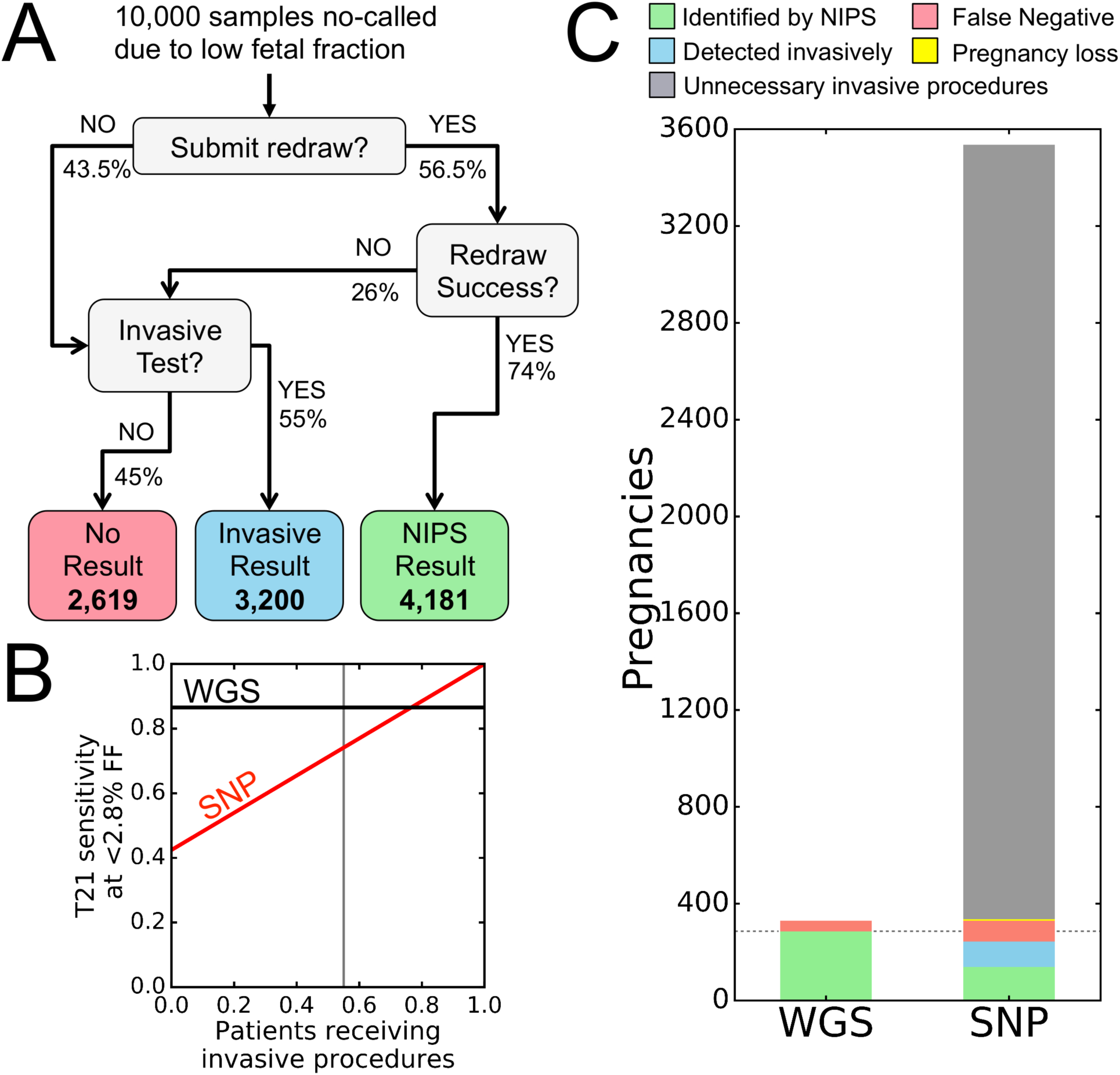
Consequences of “no-calling” samples at low fetal fraction vs. reporting at reduced sensitivity. A) Clinical decision flowchart for samples receiving a “no-call” from the SNP method. Frequencies associated with each decision branch-point were obtained from published literature (see Methods). The outcomes of 10,000 samples that receive a “no-call” result are shown in the colored boxes. B) By summing the T21 sensitivity over the frequency of fetal fractions in the range of under which the SNP method reports a “no-call”, the aggregate sensitivity of T21 detection of the WGS method is a constant 86% (black line). In contrast the SNP method (red line) is expected to detect 41% of aneuploid cases by re-draw (the intercept), while all further cases must be detected by invasive procedures. The grey line indicates a maximal estimate of the rate of patients consenting to invasive procedure given a “no-call” result (55%), which results in a total detection rate of 74%. FF, fetal fraction. C) Clinical consequences for 10,000 patients with fetal fraction below the 2.8% “no-call” threshold. While the WGS method would detect 285 out of 330 expected cases of T21 (86%), the SNP method would detect 138 by NIPS (all due to redraw). In addition, 3,200 invasive procedures would be required to detect an additional 106 cases for a net sensitivity of 74%. This would also result in six procedure-related pregnancy loss (yellow line). The dotted line indicates the number of cases of T21 detected by the WGS method for comparison.

To illustrate the clinical consequences of systematically “no-calling” low fetal fraction samples, we assessed T21 screening outcomes for both methods on a simulated cohort of 10,000 initially low fetal fraction samples, assuming a trisomy 21 incidence of 3.3% (8), and a rate of invasive testing of 55% after a “no-call” result (Figure 4C; see Methods). The WGS method would successfully detect 285 of the expected 330 cases (86%) but result in 45 false negatives. In comparison, the SNP method would detect 138 cases upon redraw, while 3,200 invasive procedures would be required to detect an additional 106 cases (74% detected), totaling 86 false negatives and six procedure-related pregnancy losses.

## Discussion

The vast majority of samples submitted for NIPS have sufficient fetal fraction to enable screening by both methods with excellent sensitivity, likely explaining why meta-analyses have noted comparable test performance (2,3,7,8). Nevertheless, we demonstrate that there are substantial performance differences between the two methodologies in low fetal fraction samples, leading to important clinical consequences.

### Modeling NIPS at low fetal fraction

We implemented a modeling approach to analyze the performance of the two NIPS methods due to a paucity of clinical data in samples with low fetal fraction. The key parameters of the WGS model - the distribution of counts along chromosomes as well as total reads per sample - have been well characterized, indicating that the results of our simulated model likely reflect biological reality for most patients (16). For the SNP model, two key parameters - the number of reads per SNP and the accuracy with which allele fractions are estimated - were inferred from published reports (see Supplemental Methods, Supplemental Figure S2) (10,25,26). Importantly, we show that even when these parameters are set to biologically unrealistic, theoretical ideals, the conclusions of the analysis remain unaltered (Supplemental Figures S3-5).

### Performance of NIPS methods

By comparing each chromosome to a baseline disomic distribution, the WGS method maintains high specificity(≥ 99.87%), independent of fetal fraction (Figure 3A) (9). In contrast, by evaluating the likelihood of aneuploidy on a sample-by-sample basis without a baseline expectation, the SNP method is unable to clearly distinguish disomic from trisomic samples at low fetal fractions. This justifies establishment of a threshold below which all samples are “no-called” (Figure 3B). In fact, the threshold of 2.8% recently reported by Ryan et al. (11) agrees well with the value below which our simulations show a substantial drop in sensitivity (i.e., 3%, Figure 3C,D).

A key determinant of the sensitivity of the SNP method in detecting trisomies is the origin of the nondisjunction event (Figure 2). The low sensitivity of detection of the most common type of nondisjunction (maternal M1, 70% of cases) overwhelms the improved sensitivity of the remaining types of nondisjunction, leading to overall poor test performance at low fetal fractions. Crucially, this performance deficit is not limited by the SNP method algorithm, but rather by the underlying biology of nondisjunction (Figure 2B, Table 1). Indeed, any method measuring shifts in allele balances within the maternal-fetal mixture of cfDNA will have relatively reduced sensitivity in samples with the most common type of nondisjunction.

We also note that the results presented here model analytical sensitivities and specificities. Reported clinical sensitivities of NIPS have been higher for T21 when compared to T18 and T13 (7,8). This is potentially due to T21 samples showing slightly elevated fetal fractions relative to disomic pregnancies, while those of T18 - and in some cases T13 – have reportedly lower fetal fractions, further illustrating the importance of maximizing detection performance in low fetal fraction samples (27,28). Other factors that reduce clinical sensitivity, such as sample contamination and confined placental mosaicism (29), should impact all NIPS methods as currently practiced (7).

### Clinical consequences of no-calling low fetal fraction samples

Our model demonstrates that while setting a minimum fetal fraction threshold may help to maintain high per-patient analytical sensitivity (e.g., 1,11,30,31), it may ultimately prove to be counter-productive. Based upon published clinical data, the probability of detecting a trisomy 21 case after an initial “no-call” using the SNP method is only ˜74% (Figure 4B). This is almost certainly an overestimate as it is unlikely that patients who receive a “no-call” due to low fetal fraction will seek confirmatory invasive testing at the same rate as those who screen positive for T21 (i.e., 10,12). In contrast, the WGS model identifies a larger fraction of T21 fetuses, all noninvasively, obviating the need for invasive testing on failed samples (Figure 4C).

A major advantage of cfDNA-based NIPS over previous screening modalities is a tenfold reduction in the false-positive rate, which has vastly reduced the number of needless invasive procedures (1,19,32). However, this benefit is undermined by tests with high failure rates, most commonly due to low fetal fraction. For example, nearly three-quarters of the projected overall test failure rate of the SNP method reported in Ryan et al. (11) was due to insufficient fetal fraction (3.8% low fetal fraction; 5.2% total failure rate).

It is also notable that test failures are routinely excluded from claimed test sensitivity in published validation reports (2,19). This is especially important given that multiple studies have noted an increased rate of aneuploidies among patients who receive a “no-call” (1,11,33). For instance, the 100% sensitivity of T21 screening reported by Pergament et al. (33) drops to 86.5% if T21 positive fetuses among “no-calls” are counted among false negatives (19). Barring an increase in the rate of invasive procedures - and the concomitant iatrogenic pregnancy loss that this entails - the findings of this study suggest that the most effective approach to improving fetal screening efforts is to implement methods that improve the performance of NIPS methods at low fetal fractions.

## Conclusion

We show that unlike the WGS approach, the SNP method cannot maintain high specificity and sensitivity at low fetal fractions, justifying its reliance on a minimal fetal fraction threshold for calling (11). Finally, using published clinical data, we find that the WGS method detects a higher proportion of common aneuploidies in low fetal fraction samples than setting a “no-call” threshold, avoiding large numbers of invasive tests and associated complications.

## 1 Supplemental Analyses

### 1.1 Dependence of the WGS method on proper normalization of bin counts

The sensititvity of the WGS method is dependent on being able to detect a shift in the distribution of bin counts on aneuploid chromosomes. The composition of certain genomic regions may cause increased variability (overdispersion) in bin counts due to features such as repetitive elements that prevent unique assignment of reads to a given location, or an unusually high or low percent guanine-cytosine bases (%GC) leading to NGS library incorporation/amplification biases (e.g., Benjamini and Speed 2012, Janevski et al. 2012). If unaddressed, the increased variance in counts can have a strong e.ect on sensitivity (Figure S1). However, it has been shown that proper normalization, involving both the removal of high-variance bins and correction for the known biases introduced by %GC, can obviate this variance and produce bin counts consistent with near-ideal Poisson expectations (Fan and Quake 2010).

**Figure S1:**
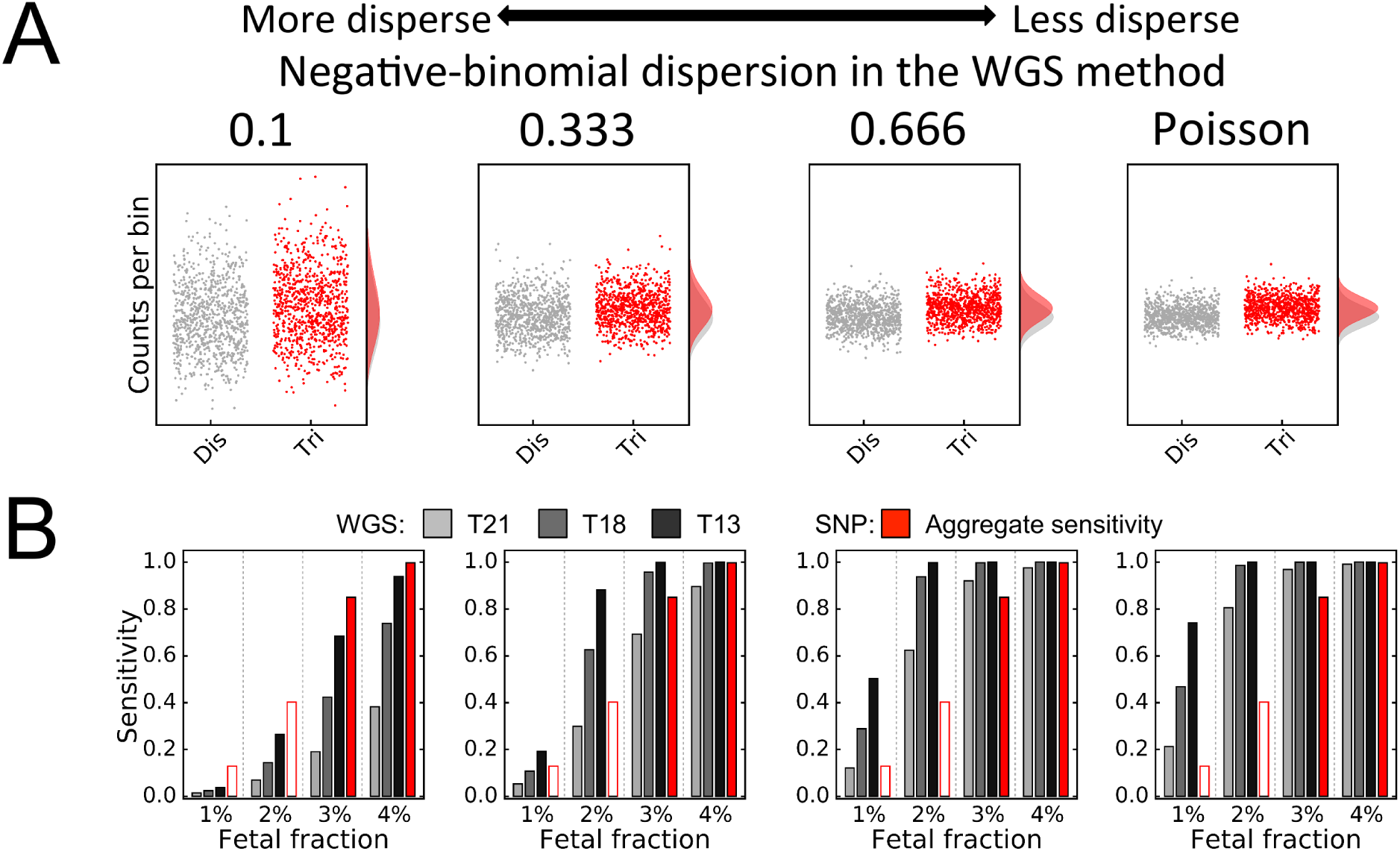
Overdispersion in bin counts relative to Poisson expectations reduces sensitivity of the WGS method. A) Plot of simulated counts-per-bin on chromosome 21 at 10% fetal fraction in either a disomic (dis) or trisomic (tri) state. Counts are sampled either from the Poisson distribution (right), or from increasingly overdispersed negative-binomial distributions where the dispersion parameter (see Supplemental Methods) is shown above each panel. The shape of the disomic (grey) and trisomic (red) distributions is shown on the right of each panel. B) Reproduction of Figure 3D, showing the negative effect of overdispersion on WGS method sensitivity for each of the common aneuploidies. The aggregate sensitivity of the SNP method is shown as hollow bars at 1% and 2% fetal fraction as these are below the threshold at which all samples are no called.

### 1.2 Fitting the parameters of the SNP model

Two key parameters of SNP method simulations were inferred by choosing values concordant with published figures: 1) the amount of dispersion in estimates of the relative frequencies of the A and B alleles and 2) the variability in total sequencing counts at each SNP (corresponding to parameters *disp_BB_* and *disp_NB_* in the Supplemental Methods below) (Figure S2).

**Figure S2:**
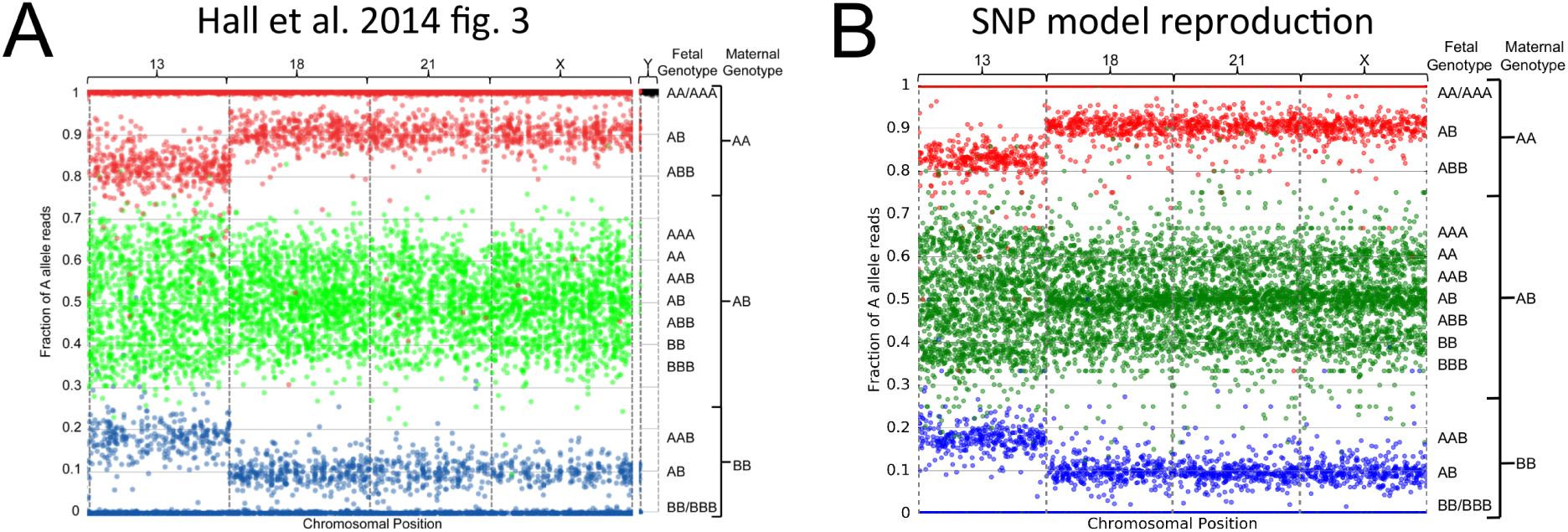
Comparison of published data from SNP method validation report (A) to the output of the SNP model using the same fetal fraction and genotypes (B). A) Figure 3 from Hall et al. (2014) featuring the output of the SNP method on a T13 fetal sample with 19.2% fetal fraction is shown unmodified. B) The output of the SNP model simulating the same chromosomal genotypes and fetal fraction reproduces the pattern observed in actual data. In fact, the simulated data (B) may be slightly under-dispersed relative to the true data (A) as the expected genotypic medians are clearly visible in the former (concentrations of points along lines). Therefore, this suggests that the model paramters are conservative and represent an upper-bound on SNP-method performance. Note that the simulated data assigns the same number of SNPs to each chromosome unlike the true data, whose chromosomal widths vary. Also, the SNP model ignored the small number of SNPs on chromosome Y, as they have no effect on this analysis.

We also note that it is likely that the simulations overestimate the performance of the SNP model because some key complications known to occur in actual samples were excluded from our analysis. First, recombination within parental chromosomes - expected in >75% of cases of maternal chromosome 21 (Kong et al. 2002) - will lead to trisomies wherein some fraction of SNPs are consistent with M1 nondisjunction, while the remainder are consistent with M2. This requires the aneuploidy caller to consider more possible hypotheses, potentially decreasing sensitivity (Rabinowitz et al. 2014). Second, fetal fraction is an explicit parameter of the calling method, as it is used to establish the expected distribution of allele fractions under the possible ploidy hypotheses. We have assumed that fetal fraction has been estimated without error; however, in reality, the accuracy of the estimate of fetal fraction will vary based on a number of parameters, including sample depth and the number of informative SNPs (Jiang et al. 2012).

Below, we further show that varying the specific values of the parameters of the SNP model do not affect the conclusions drawn from our simulations.

#### 1.2.1 Varying the dispersion in estimates of the relative frequencies of the A and B alleles, *disp_BB_*

Under ideal conditions, next-generation sequencing methods would sample reads from each allele according to binomial expectations, with mean equal to the fraction of A allele reads. In real sequencing applications various factors contribute to overdispersion of sampled allele frequencies, such as allele-specific PCR amplification bias and PCR ‘jackpot’ mutations during library preparation (Skelly et al. 2011). The degree to which allele frequencies at individual SNPs are overdispersed relative to binomial expectations is captured by the dispersion parameter of the betabinomial distribution, *disp_BB_* (see Supplemental Methods).

At low values of *disp_BB_,* the sampling accuracy of the A allele fraction decreases, making it more difficult to resolve the expected distributions of the different ploidy hypotheses (Figure S3). In contrast, as *disp_BB_* increases, the expected distributions of the different ploidy hypotheses become more distinct, thereby increasing sensitivity. In particular, as *disp_BB_* approaches infinity, the betabinomial distribution collapses to a binomial distribution, representing the upper-limit of expected sampling accuracy.

As shown in Figure S3, even under the biologically unrealistic condition of perfect binomial sampling, the fundamental conclusions of the analysis remain unaltered: the performance of the SNP method improves, but does not increase beyond that of the WGS method on any of the three common aneuploid chromosomes. Given that binomial sampling is biologically implausible (Skelly et al. 2011), the value of *disp_BB_* chosen in the manuscript, 1,000, likely better reflects the true performance of the SNP test.

**Figure S3:**
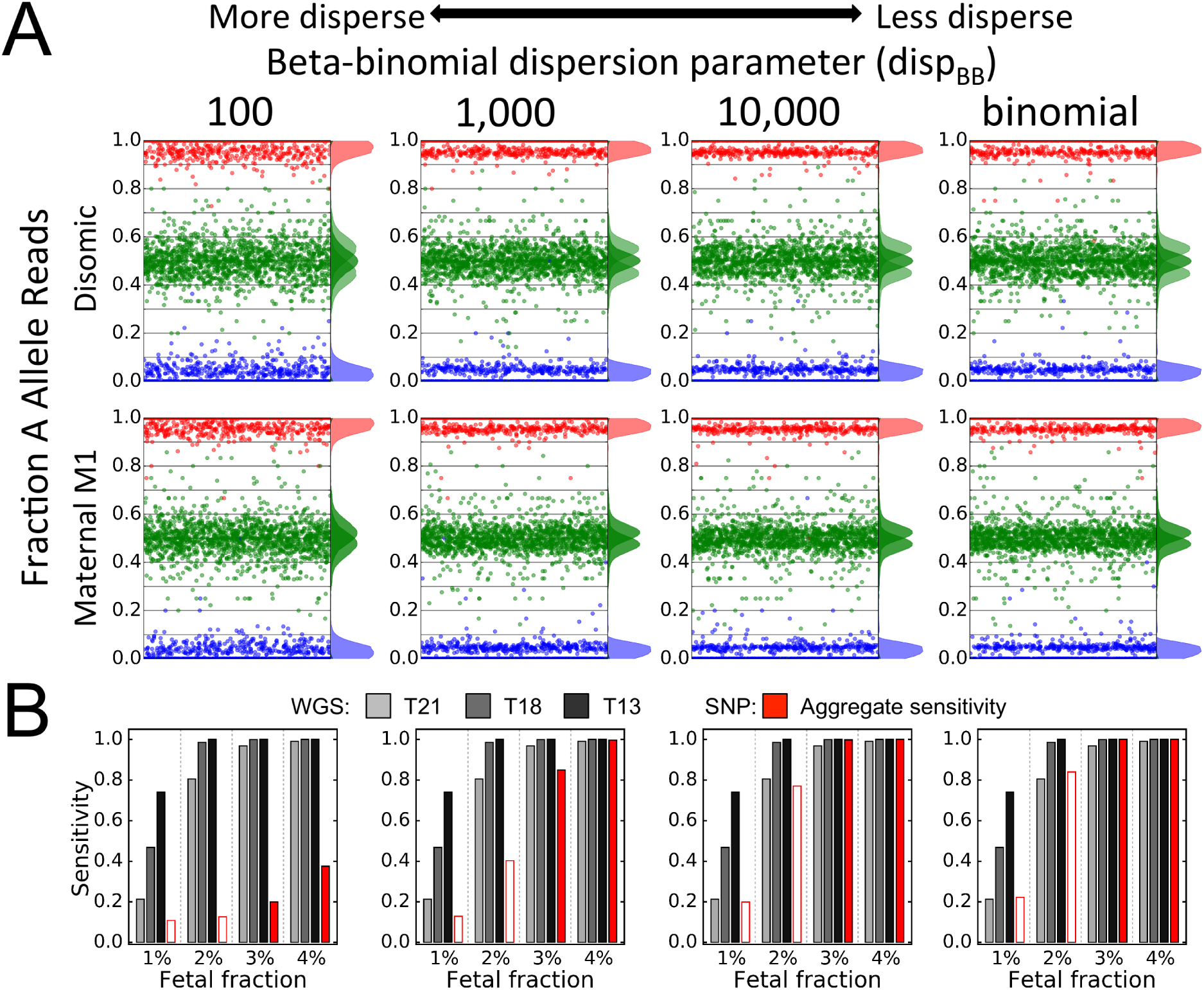
Illustration of the effect of varying the dispersion in estimates of the relative frequencies of the A and B alleles, *disp_BB_*. A) The accuracy of measurements of the expected frequency of the A allele increase with increasing values of *disp*_BB_, as shown by the width of the distributions plotted to the right of simulated diploid (top) and maternal M1 aneuploid (bottom) samples. Simulated samples are shown at 10% fetal fraction. B) Reproduction of Figure 3D, illustrating the effect of varying the beta-binomial dispersion parameter on SNP method aggregate sensitivity. Higher values of *disp_BB_* increase the SNP method caller’s ability to discriminate between ploidy hypotheses, thus increasing sensitivity at a given fetal fraction. As *disp_BB_* approaches infinity, sampling becomes binomially distributed (rightmost panels), representing the limit of expected sampling accuracy. Even in cases of ideal sampling, performance of the SNP method remains below that of the WGS method on any chromosome. Note that 1,000 is the value of *disp_BB_* used for the simulations in the main manuscript. The aggregate sensitivity of the SNP method is shown as hollow bars at 1% and 2% fetal fraction as these are below the threshold at which all samples are no called.

#### 1.2.2 Varying the variability in total sequencing counts at each SNP, *disp_NB_*

Due imperfect sampling of SNPs during sequencing during sequencing library preparation, counts-per-SNP are overdispersed relative to Poisson expectations (Anders and Huber 2010). This overdispersion is captured by the dispersion parameter of a negative-binomial distribution, where 0 < *disp_NB_* < 1 (see Supplemental Methods).

At values approaching 1, *disp_NB_* collapses to the Poisson distribution, and thus ideal sampling. As *disp_NB_* approaches 0, the variability in counts-per-SNP increases, leading to some SNPs being under-sampled and non-informative, while others are sampled at much higher depth. However, the SNP caller explicitly takes into account total counts in its assignment of likelihood at each SNP (see Supplemental Methods). Since the number of SNPs interrogated at each chromosome is large, the reduction in confidence at low depth SNPs is offset by the increased confidence at high-depth SNPs, and therefore the performance of the caller is largely independent of the specific value chosen for *disp_NB_* (Figure S4)

**Figure S4:**
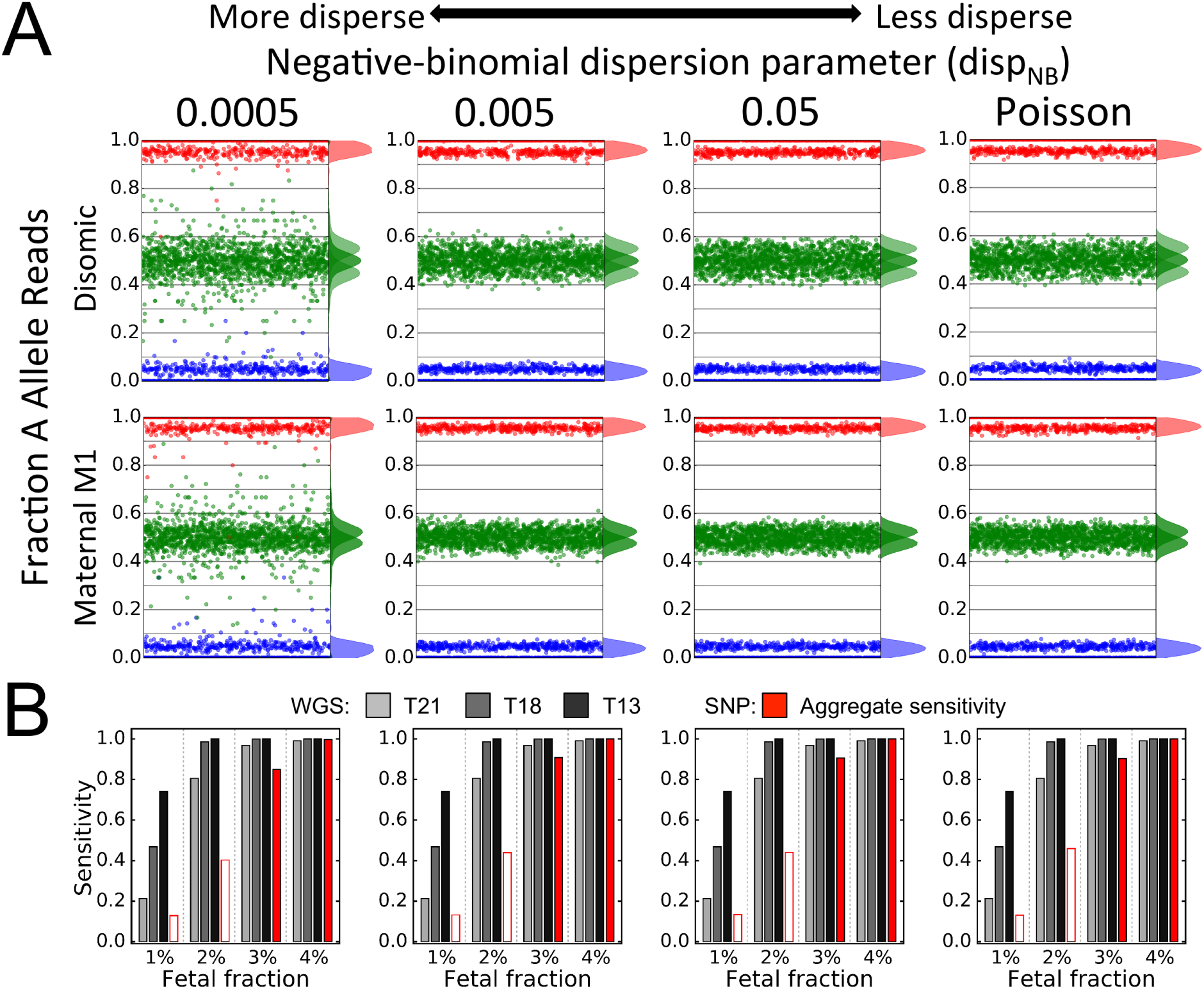
Illustration of the effect of varying the dispersion in total sequencing counts at each SNP, *disp_NB_*. A) Decreasing values of *disp_NB_* increases the number of SNPs with low sequencing counts, thereby increasing stochastic sampling ‘noise’ as shown by SNPs falling far from the centers of the expected frequency distributions of simulated diploid (top) and maternal M1 aneuploid (bottom) samples. Simulated samples are shown at 10% fetal fraction. As the value of *disp_NB_* approaches 1, sampling approaches Poisson expectations (rightmost panels). B) Reproduction of Figure 3D, illustrating the effect of varying the negative-binomial dispersion parameter on SNP method aggregate sensitivity. As can be seen, varying of this parameter has almost no effect on the SNP method caller’s performance, even when sampling under biologically implausible, idealized Poisson parameters. Note that the leftmost value of 0.0005 is that used for the simulations in the main manuscript. The aggregate sensitivity of the SNP method is shown as hollow bars at 1% and 2% fetal fraction as these are below the threshold at which all samples are no called.

#### 1.2.3 Varying the number of SNPs interrogated on a chromosome

The overall likelihood that a chromosome is consistent with a particular ploidy hypothesis is the sum of the individual likelihoods calculated at each SNP (Rabinowitz et al. 2014). For the purposes of the analyses reported in the main manuscript, we have assumed that the 13,392 SNP sites were evenly distributed across the four interrogated chromosomes (i.e., 3,348 SNPs on each of chromosomes 21, 18, 13, and X). Nevertheless, we assessed the effect on sensitivity of lowering or increasing the number of interrogated SNPs on a single chromosome.

As can be seen in Figure S5, reducing the number of SNPs causes a substantial decrease in sensitivity at all fetal fractions. Conversely, increasing the number of SNPs leads to only to modest gains in sensitivity: even interrogating 10,000 SNPs on a single chromomosome does not raise the sensitivity of the SNP method to the performance to the WGS method on chromosome 21, and therefore does not a?ect theconclusions ofour analysis.

**Figure S5:**
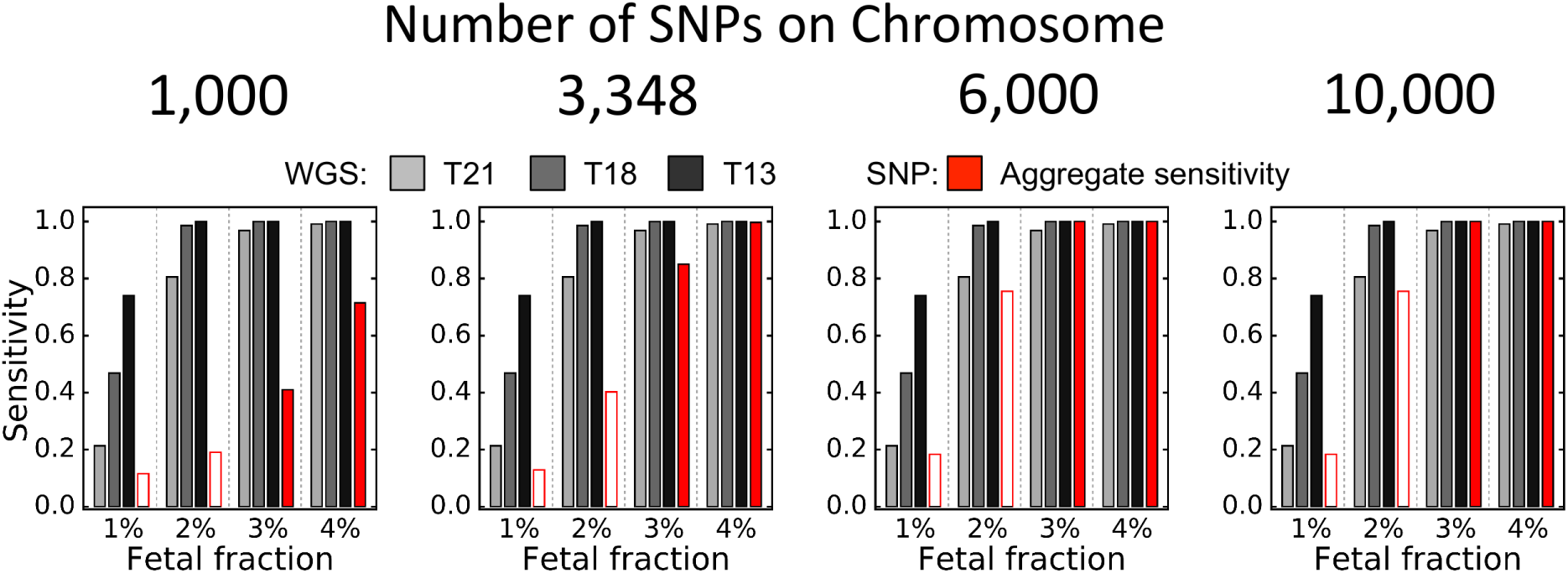
Reproduction of Figure 3D illustrating the effect of varying the number of SNPs interrogated on a single chromosome. Reducing the number of SNPs decreases aggregate sensitivity of the SNP method across all fetal fractions, whereas increasing the number of SNPs leads to relatively modest gains. The aggregate sensitivity of the SNP method is shown as hollow bars at 1% and 2% fetal fraction as these are below the threshold at which all samples are no called.

### 1.3 Overview of informative genotypes under the four origins of nondisjunction

**Figure S6:**
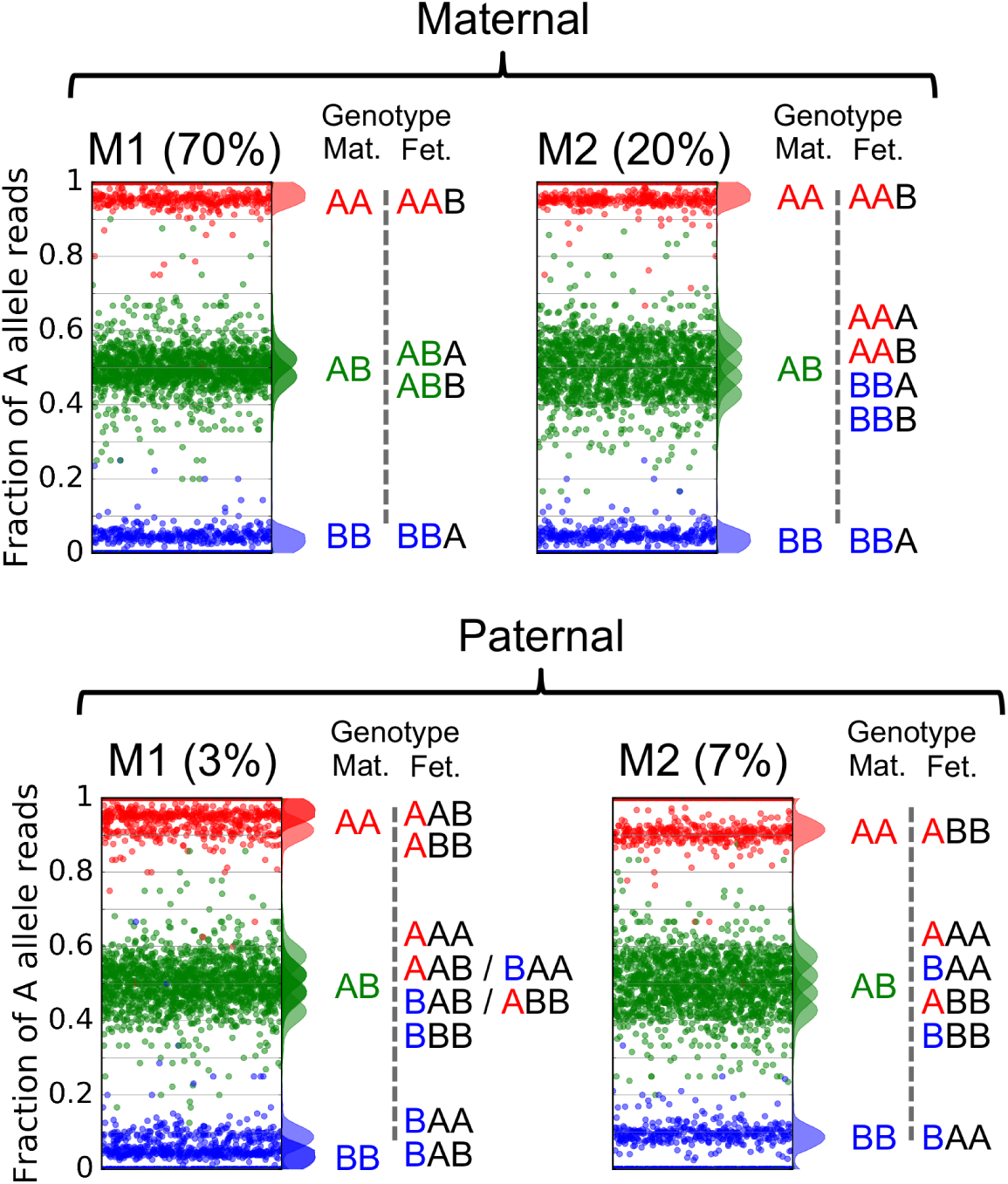
Possible informative maternal-fetal genotype combinations under the four origins of nondisjunction. The four possible origins of nondisjunction are shown along with the expected A allele fraction distributions from possible informative maternal-fetal genotype combinations (e.g., 100% A or B allele distributions are not considered). Maternal genotypes are color-coded: AA, red; AB, green; BB, blue. The alleles from maternally inherited fetal chromosome(s) are colored the same, with a single A allele also colored red and a single B allele colored blue. The alleles from paternally inherited chromosome(s) are colored black. As can be seen, each origin of nondisjunction produces a unique combination of genotypes, which form the basis of the SNP-method test.

### 1.4 Estimating the distribution of fetal fractions

The distribution of fetal fractions was obtained by fitting a beta distribution to the paramaters indicated in Nicolaides et al. (2012): a median of 0.10 with interquartile range 0.078 to 0.13 (see Methods in main manuscript). Fit parameters were a = 6.524 and β = 55.606 (Figure S7).

**Figure S7:**
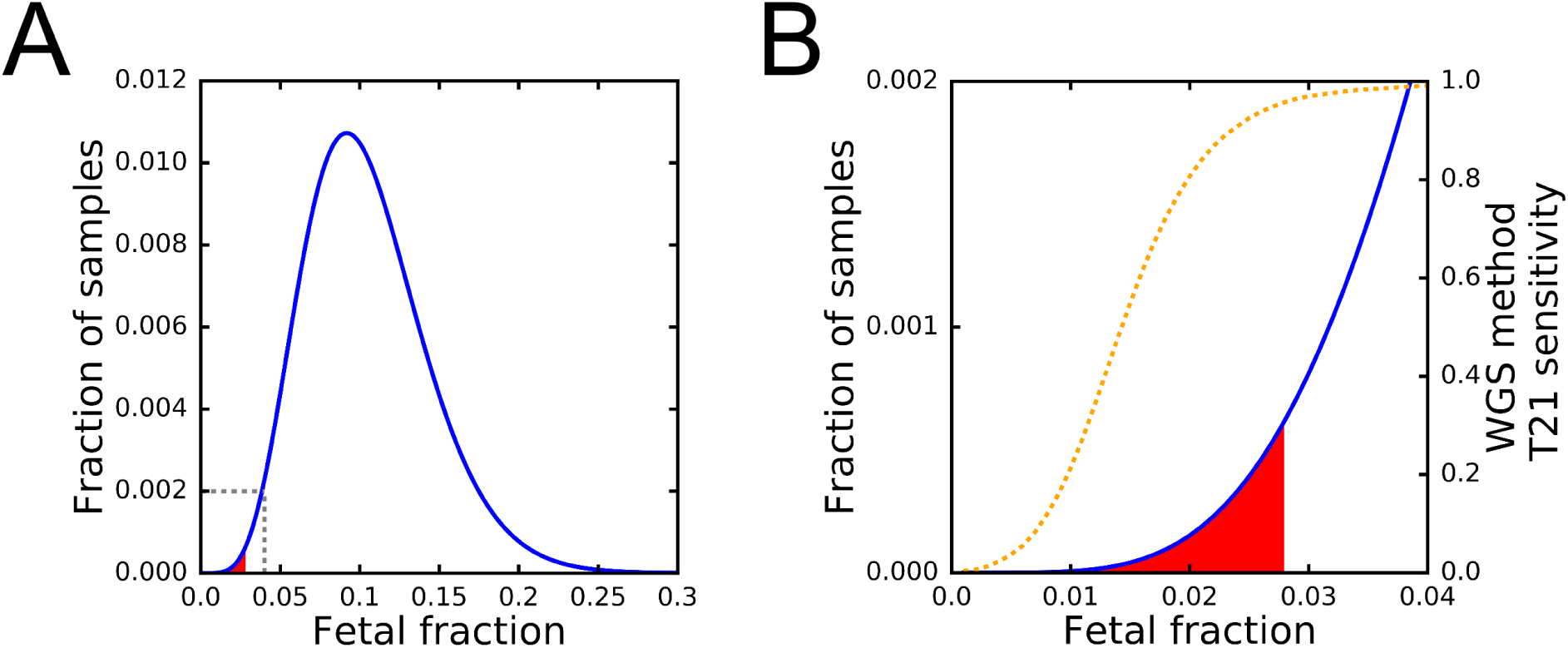
Fetal-fraction distribution inferred from Nicolaides et al. (2012) A) Distribution of fetal fractions fit to data of Nicolaides et al. (2012). The shaded red area indicates samples that would be no called below a threshold fetal fraction of 2.8% (Ryan et al. 2016). The region bounded by the dashed grey lines is enlarged in panel B. B) The same distribution is shown, focusing on the region < 4% fetal fraction. The dashed orange line is the sensitivity of T21 detection of the WGS method. The sensitivity values were estimated in 0.1% fetal fraction increments (see Methods), and a second-order, Savitzky-Golay filter with a step of seven datapoints was applied to interpolate continuous values (Savitzky and Golay 1964).

There exists some systematic discrepancy in how different methods estimate fetal fraction. Based on the distribution in Figure S7, 0.33% of samples will fall below the indicated 2.8% fetal fraction threshold. This is in-line with Porreco et al. (2014) and Lefkowitz et al. (2016) who reported that 1% of samples fell below their 4% fetal fraction threshold (according to our distribution, 1.9% of samples will have less than 4% fetal fraction). In contrast, projection Ryan et al.’s’ (2016) validation sample fetal-fraction distribution to that expected from clinical samples, the authors estimate that 3.8% of samples will fall below their 2.8% fetal fraction threshold.

## 2 Supplemental Methods

### 2.1 WGS method simulations

#### 2.1.1 Simulating WGS method samples

The WGS method compares the distribution in the number of reads mapping to fixed-sized bins across a chromosome to the background distribution of all samples sequenced on the same sequencing flowcell (hereafter, ‘batch’), in order to determine whether the counts derived from the sample in question deviate significantly from expectations.

The lengths of each of the interrogated chromosomes were obtained from the GRCh37/hg19 (Feb. 2009) release of the human genome (Lander et al. 2001) and divided into bins of 50 kb. In order to account for the removal of poorly mappable and high GC bins (Jensen et al. 2013), 10% of the bins on each chromosome were discarded (Table S1).

**Table S1:** Chromosomal lengths and number of bins according to human genome release hg19.

| Chrom | Length | Bins |
| --- | --- | --- |
| 13 | 115,169,878 | 2,073 |
| 18 | 78,077,248 | 1,405 |
| 21 | 48,129,895 | 866 |

As noted in Fan and Quake (2010), when normalized for GC content and mappability, counts of reads mapping to each bin along a given chromosome follow the expectations of a Poisson distribution. Batches of 100 samples for each interrogated chromosome were generated at random by sampling from a population with trisomic fetuses at the following frequencies: T21, 3.3%; T18, 1.5%; and T13, 0.5% (Taylor-Phillips et al. 2016). The mean counts-per-bin, 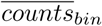, for each sample was obtained from Jensen et al. (2013) where:

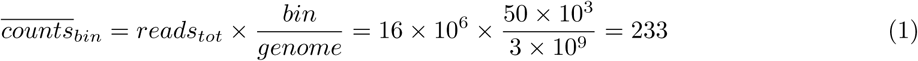

For healthy diploid samples, counts-per-bin for the number of bins corresponding to the chromosome of interest were drawn from a Poisson distribution with mean 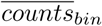. For aneuploid samples, bin counts were drawn from the same distribution with the mean multiplied by 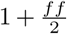 in order to account for the additional counts originating from the duplicated fetal chromosome, where *ff* is the fetal fraction of non-aneuploid chromosomes.

#### 2.1.2 Calling trisomies via the WGS method

The distribution of counts-per-bin for each sample was used to calculate a z-score:

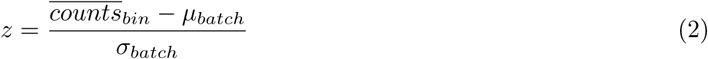

where 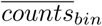 is the mean counts-per-bin of the sample, and *μ_batch_* and *σ_batch_* are the mean and standard deviation of the mean counts-per-bin of the other samples in the batch (i.e., not including the sample whose z-score is being calculated), respectively.

Samples with z ≥ 3 were called trisomic for the chromosome of interest.

#### 2.1.3 Calculating sensitivity and specificity of the WGS method

For each of the interrogated chromosomes, batches were simulated at fetal fractions 0.001 to 0.04 in increments of 0.001. All simulated fetal fractions from 0.001 to 0.027 were used to calculate the aggregate sensitivity of the WGS method for all samples in this range of fetal fractions (see Main Methods), while batches simulated at fetal fractions 0.01, 0.02, 0.03, and 0.04 were used to plot Figure 3. From each batch, one diploid and one aneuploid z-score (if present) were sampled at random until 10,000 of each category were sampled. Sensitivity and specificity were then calculated as:

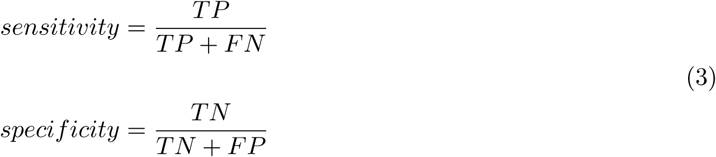

where *TP* and *FN* are the counts of aneuploid samples correctly called aneuploid or incorrectly called diploid, respectively, and *TN* and *FP* are the counts of the diploid samples correctly called diploid or falsely called aneuploid, respectively.

### 2.2 SNP method simulations

#### 2.2.1 Simulating SNP method samples

Samples used to interrogate the properties of the SNP method were simulated as follows. For each sample, corresponding to a single chromsosome, we simulated 3,348 SNP sites, representing an even distribution of the 13,392 primer pairs reported in Ryan et al. (2016) across the four interrogated chromosomes (21, 18, 13, and X). For the purposes of our simulations, we assumed that SNPs along the chromosome showed no linkage disequilibrium and that no recombination took place between parental chromosomes. We note however, that incorporation of both of these parameters would act to reduce the sensitivity of the method (Rabinowitz et al. 2014). Phased maternal and paternal genotypes (AA, AB, and BB) along the chromosome were randomly generated assuming Hardy-Weinberg equilibrium (Equation 4) and population frequency of the A allele, *f_A_*, was drawn from a beta distribution with mean 0.5 and standard deviation 0.1.

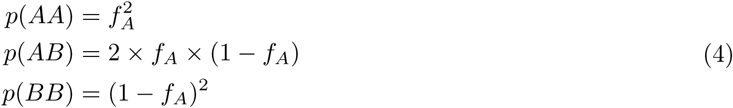

Simulated parental chromosomes were then randomly segregated to create a fetal genotype according to the specified chromosomal state (diploid, maternal M1 aneuploidy, maternal M2 aneuploidy, paternal M1 aneuploidy, or paternal M2 aneuploidy).

The mean read depth, and thus sum of the counts of both alleles, was set to 859, corresponding to a mean sequencing depth of 11.5 × 10^6^ reads, distributed over the 13,392 sites as reported in Ryan et al. (2016). It is known that variability in NGS read depth is well-captured by a negative binomial model, *NB*(*r*,*p*), (Anders and Huber 2010). Given the constraint that mean of the distribution must be equal to the mean read depth, it is possible to express the distribution in terms of a single dispersion parameter, *disp_NB_*:

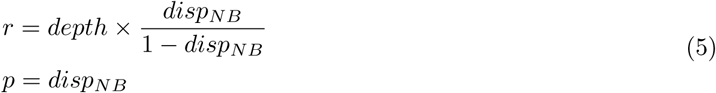

As the degree of dispersion in read depth is not reported, *disp_NB_* was simulated over a range of values and 0.0005 was chosen as it produced data concordant with published figures (Figure S2).

If NGS library preparation perfectly captured the relative allelic proportions in the circulating maternal blood and allelic sampling were perfect during sequencing, we would expect the number of A allele reads at each site to be binomially distributed, *B(p(A), depth)*, with probability of sampling the A allele equal,*p*(*A*):

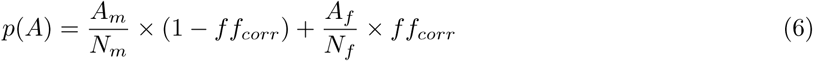

where *A_m_*, and *A_f_* are the number of A-containing chromosomes in the maternal and fetal genotypes, respectively, while *N_m_* and *N_f_* are the total number of maternal and fetal chromosomes, respectively. In the case of chromosomal aneuploidy, the chromosome-specific fetal fraction will be different than that estimated from the overall genome. The chromosome-specific ‘corrected’ fetal fraction, *ff_corr_*, is obtained from:

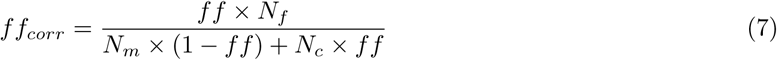

However, it is clear that published data are overdispersed relative to binomial expectations (Figure S2) (Skelly et al. 2011). Such overdispersion is well-captured by the beta-binomial distribution, *betabin*(*n*, *α*,*β*), which allows *p*(*A*) to be innacurately represented in sequence counts according to a set amount of dispersion. Again, given the constraint that dispersion in *p*(*A*) be centered around *p*(*A*) itself, it is possible to represent the dispersion parameters, *α* and *β*, in terms of a single dispersion parameter *disp_BB_*:

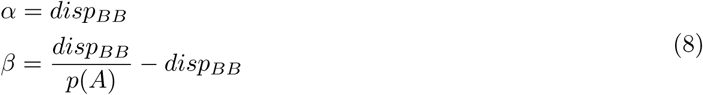

As above, the degree of dispersion in allele sampling is not published, and it was necessary to simulate data over a range of parameters in order to determine a value of *disp_BB_* consistent with published data (Hall et al. 2014). The value used in the main manuscript was 1,000; however, we show that varying this parameter does not alter the conclusions of this study (see Supplemental Analysis).

For each simulated sample, at each of the 3,348 SNP sites, the maternal genotype, total read depth, number of A reads, and population frequency of the A allele were recorded and used to test the calling performance of the SNP method.

### 2.2.2 Calling trisomies via the SNP method

The method described here is adapted from the description in Rabinowitz et al. (2014). Under the SNP method, our goal is to test which of the possible chromosomal ploidy hypotheses (diploid, maternal M1 or M2, and paternal M1 or M2) is most consistent with the observed data:

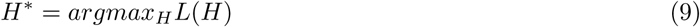

where *H*\* is the most likely hypothesis, and *L*(*H*) is the likelihood of the hypothesis being correct, given the data.

The data that we observe are counts at individual SNPs. Therefore, assuming no recombination, the chromosome-level *L*(*H*) is simply the sum of the likelihoods calculated for each individual SNP, *L*(*i,H*):

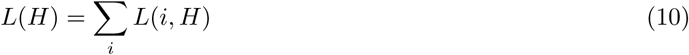

At the SNP-level, we can calculate the probability of observing a particular allelic ratio, *x,* given the maternal genotype, *gm*, the fetal genotype, *gf*, the fetal fraction *ff*, and the chromosomal ploidy hypothesis, *H*, or *p*(*x|gm, gf, ff*, *H*). As the fetal genotype, *gf*, is not known, we must marginalize over the probability of each fetal genotype:

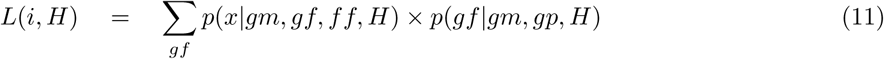

where *gp* is the paternal genotype. Note that here we assume that the true fetal fraction is known.

### 2.2.3 Calculating *p*(*gf*|*gm*, *gp*, *H*)

Given knowledge of both parent’s genotypes, *p*(*gf*|*gm*, *gp*, *H*) is easily calculated. For example, under the hypothesis of diploidy, if *gm* is AA and *gp* is AB, then the probability that *gf* is AA is equal to 0.5, and so on. However, as the SNP method is currently implemented, the paternal genotype is not determined independently (Ryan et al. 2016). Consequently, it is necessary to sum over the individual probabilities of each possible paternal genotype.

If the population frequency of the segregating alleles is known, then it is possible to infer the probability of each possible paternal genotype, *p*(*gp*), assuming that the SNPs being interrogated are in Hardy-Weinberg equilibrium. The probability of the fetal genotype can then calculated by summing over possible paternal genotypes weighed by their population probability:

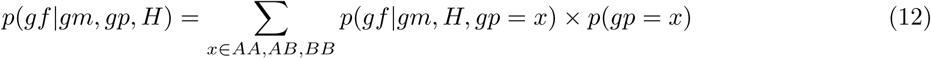

where *p*(*x*) is obtained from equation 4.

### 2.2.4 Calculating *p*(*x*|*gm*, *gf*, *ff*, *H*)

In order to determine the probability of observing the data, *p*(*x*|*gm*, *gf*, *ff*, *H*), the counts of the A allele at the locus, it is first necessary to determine the probability of sampling a read containing the A allele, *p*(*A*), according to equation 6.

Assuming that allele counts are sampled from a beta-binomial distribution (see simulating data above), we can obtain the probability of observing the data from the distribution itself:

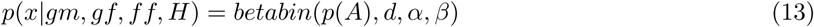

where *d* is the total counts at the SNP, and *α* and *β* are the parameters of the beta-binomial distribution, which are known from the simulation parameters (equation 8).

### 2.2.5 Assigning a log-odds ratio to the likelihood of aneuploidy

Once the likelihood of each individual chromosomal ploidy hypothesis has been determined, a log-odds ratio of the likelihood of diploidy, *LOR,* was determined as:

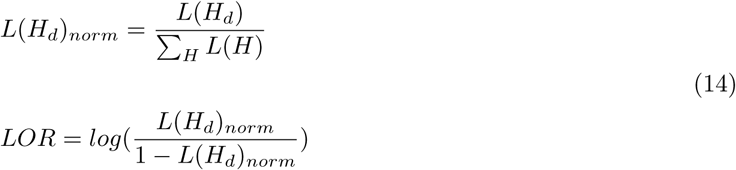

where *L(H_d_)_norm_* is the fraction of the total likelihood represented by the hypothesis of diploidy. Positive *LOR* values indicate chromosomes with a higher likelihood of diploidy, while negative values indicate chromosomes with a higher likelihood of any trisomy.

## 2.3 Generating ROC Curves

In order to generate ROC curves representing the aggregate sensitivity of the SNP method, at each fetal fraction (0.1-4% in 0.1% increments), we randomly sampled 1,000 LORs from the following ploidy hypotheses at the specified frequency: M1 maternal trisomy (70%), M2 maternal trisomy (20%), M1 paternal trisomy (3%), and M2 paternal trisomy (7%). We also sampled 1,000 random disomic fetus LORs and calculated AUCs using the `metrics.auc()` function in the sklearn package in Python (Pedregosa et al. 2011), version 0.17. The reported mean AUCs and their 95% confidence intervals were calculated from 1,000 permutations of the above sampling approach.

AUCs for T21, T18, and T13 under the WGS method were generated similarly in order to provide directly comparable values: At each fetal fraction, we randomly sampled 1,000 z-scores from aneuploid chromosomes as well as 1,000 z-scores from diploid chromosomes. As above, AUCs and their 95% confidence intervals were calculated from 1,000 permutations of this sampling.

